## supplemental for "Ligand bias underlies differential signaling of multiple FGFs via FGFR1"

### Scaling of phosphorylation western blot data

To identify and quantify bias, the phosphorylation dose response curves for the three ligands have to be globally scaled such that the efficacy of the full agonist is set to 1, and the efficacies of the partial agonist are scaled accordingly. Below we present the protocol. Steps 1 through 3 are used in the literature to scale the individual dose response curves. Steps 4 through 6 implement the global scaling.

1. Collect at least three independent dose response curves for each ligand. Each dose response is acquired using one gel.

2. Scale each dose response curve so max is 1. In other words, set the band intensity for the sample with the highest band intensity to 1.

3. Average the dose response curves, and reset the maximum value to 1 if needed.

4. For each ligand, re-run 3 of the samples with intensity 1 on a gel. This is the so-called "gluing gel". This gel includes 3 samples for each ligand, so 9 total. Example of such a gluing gel for pFRS2 is shown here, showing 3 independent samples for fgf4 (left), fgf8 (middle), and fgf9 (right)

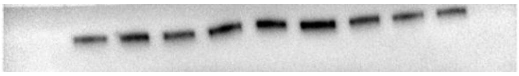

5.  Average the three intensities for each ligand. Identify the ligand that yielded the highest average intensity. Rescale the three averages, such that this highest intensity is set to 1.  For the case of pFRS2 shown above, the scaled averages are 0.81 ± 0.09, 1.00±0.03 and 0.51 ± 0.10. These are the global scaling coefficients.

6. Multiply the averaged dose response curves by the respective global scaling coefficients to obtain the globally scaled dose response curves.

**Scaling of FGFR1 and collagen abundance gels**

In this case, the band intensities at zero ligand are the highest and they are set to 1; the rest are scaled accordingly. Results from at least 3 independent samples are averaged. Then, 1 is subtracted from all the data to obtain loss of abundance dose responses.

Table S1: $\beta'_4$, $\beta'_8$, and $\beta'_9$ values calculated for each response using equation 4

| FGF4 | | | | | |
|---|---|---|---|---|---|
| | pY653/654 | pY766 | pPLCγ | pFRS2 | Downregulation |
| pY653/654 | | 0.185 ± 0.096 | -0.702 ± 0.112 | -0.140 ± 0.093 | 0.734 ± 0.218 |
| pY766 | | | -0.887 ± 0.144 | -0.325 ± 0.132 | 0.549 ± 0.231 |
| pPLCγ | | | | 0.562 ± 0.155 | 1.435 ± 0.241 |
| pFRS2 | | | | | 0.873 ± 0.229 |
| Downregulation | | | | | |
| FGF8 | | | | | |
| | pY653/654 | pY766 | pPLCγ | pFRS2 | Downregulation |
| pY653/654 | | 0.116 ± 0.129 | -0.946 ± 0.135 | -0.753 ± 0.128 | 0.372 ± 0.150 |
| pY766 | | | -1.062 ± 0.118 | -0.870 ± 0.110 | 0.256 ± 0.136 |
| pPLCγ | | | | 0.193 ± 0.126 | 1.318 ± 0.148 |
| pFRS2 | | | | | 1.125 ± 0.136 |
| Downregulation | | | | | |
| FGF9 | | | | | |
| | pY653/654 | pY766 | pPLCγ | pFRS2 | Downregulation |
| pY653/654 | | 0.065 ± 0.152 | -0.956 ± 0.102 | 0.026 ± 0.079 | 0.734 ± 0.132 |
| pY766 | | | -1.020 ± 0.209 | -0.038 ± 0.202 | 0.670 ± 0.220 |
| pPLCγ | | | | 0.982 ± 0.101 | 1.690 ± 0.145 |
| pFRS2 | | | | | 0.708 ± 0.123 |
| Downregulation | | | | | |

Table S2: Comparison of β'$_4$, β'$_8$, and β'$_9$ values from Table S1. Shown are p values calculated by ANOVA. P values < 0.5 are highlighted in gray.

| β' Significance Values | | | |
|---|---|---|---|
| | 4v8 | 4v9 | 8v9 |
| pY653/654 vs pY766 | >0.05 | >0.05 | >0.05 |
| pY653/654 vs pPLCγ | >0.05 | >0.05 | >0.05 |
| pY653/654 vs pFRS2 | 0.001 | >0.05 | 0.0002 |
| pY766 vs pPLCγ | >0.05 | >0.05 | >0.05 |
| pY766 vs pFRS2 | 0.025 | >0.05 | 0.0015 |
| pPLCγ vs pFRS2 | >0.05 | >0.05 | 0.0014 |
| pY653/654 vs DownRegulation | >0.05 | >0.05 | >0.05 |
| pY766 vs DownRegulation | >0.05 | >0.05 | >0.05 |
| pPLCγ vs DownRegulation | >0.05 | >0.05 | >0.05 |
| pFRS2 vs DownRegulation | >0.05 | >0.05 | >0.05 |
| Collagen Loss vs Growth Arrest | 0.0002 | 0.015 | 0.0007 |

Table S3: Best-fit Gaussian Parameters for the different log(brightness) distributions

| Gaussian Fit Parameters | | |
|---|---|---|
| | mean | stdev |
| FGFR1 + 130 nM FGF9 | 0.41 ± 0.01 | 0.30 |
| FGFR1 + 130 nM FGF4 | 0.48 ± 0.01 | 0.31 |
| FGFR1 + 3 nM FGF4 | 0.30 ± 0.01 | 0.22 |
| FGFR1 + 130 nM FGF8 | 0.40 ± 0.01 | 0.25 |
| LAT | 0.22 ± 0.01 | 0.32 |
| TrkA + 130nM NT3 | 0.39 ± 0.01 | 0.29 |
| FGFR1 no ligand | 0.25 ± 0.01 | 0.31 |

Table S4: Calculated Z values from the Gaussian fit parameters in Table S3.

| Z values for Gaussians | | | | | | |
|---|---|---|---|---|---|---|
| | FGFR1 + 130nM FGF4 | FGFR1 + 3nM FGF4 | FGFR1 + 130nM FGF8 | LAT | TrkA + 130nM NT3 | FGFR1 no ligand |
| FGFR1 + 130nM FGF9 | 2.664092 | 4.477257 | 0.318121 | 5.901904 | 0.436884 | 4.868914751 |
| FGFR1 + 130nM FGF4 | | 10.4334 | 3.379447 | 9.628828 | 3.421783 | 7.801433959 |
| FGFR1 + 3nM FGF4 | | | 4.768932 | 3.348705 | 3.221729 | 3.221729253 |
| FGFR1 + 130nM FGF8 | | | | 6.089074 | 0.138045 | 4.910927558 |
| LAT | | | | | 5.825875 | 0.37353812 |
| TrkA+130nM NT3 | | | | | | 4.24156545 |

Table S5: Summary of results shown in Figures 5A-C.

| | Endocytosis fluorescence | Apoptosis slope | Viability slope |
|---|---|---|---|
| FGF8 | 0.75 ±0.04 | 0.0067 ± 0.0008 | -0.0001 ± 0.0001 |
| FGF9 | 0.94 ±0.05 | 0.0032 ± 0.0011 | -0.0017 ± 0.0003 |
| No ligand | 1.00 ±0.04 | | |

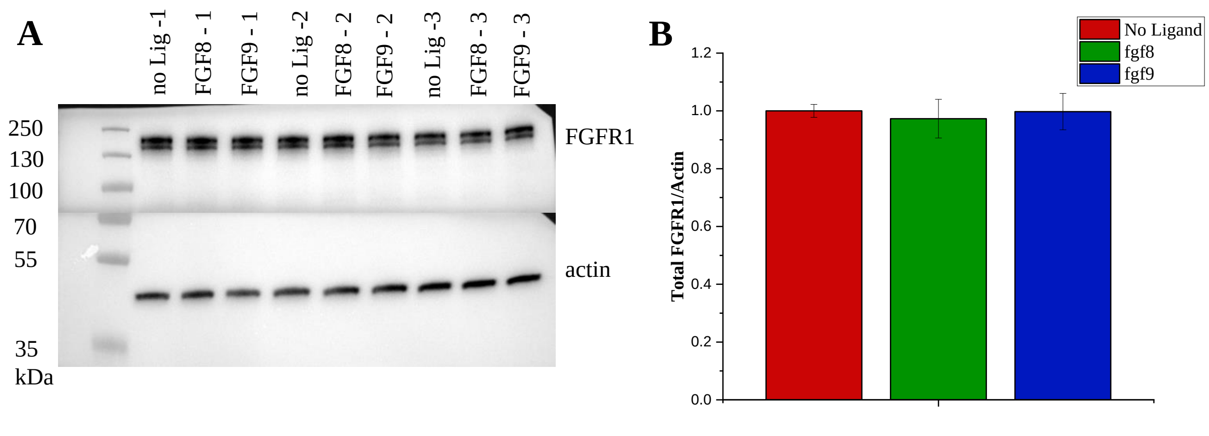


Figure S1. Total cellular expression of FGFR1 in the stable FGFR1 line. Shown are results for three different samples in the absence of ligand, three samples that have been treated with FGF8 for 2 mins, and three samples have been treated with FGF9 for 2 mins. All the expressions are the same.
